# Genomic screening of antimicrobial resistance markers in UK and US *Campylobacter* isolates highlights stability of resistance over an 18 year period

**DOI:** 10.1101/2021.08.24.457564

**Authors:** Arnoud H.M. van Vliet, Siddhartha Thakur, Joaquin M. Prada, Jai W. Mehat, Roberto M. La Ragione

**Affiliations:** School of Veterinary Medicine, Department of Pathology and Infectious Diseases, Faculty of Health and Medical Sciences, University of Surrey, Guildford GU2 7AL, United Kingdom; School of Veterinary Medicine, Department of Veterinary Epidemiology and Public Health, Faculty of Health and Medical Sciences, University of Surrey, Guildford GU2 7AL, United Kingdom; School of Biosciences and Medicine, Faculty of Health and Medical Sciences, University of Surrey, Guildford GU2 7XH, United Kingdom; Department of Population Health & Pathobiology, College of Veterinary Medicine, North Carolina State University, Raleigh, NC 27607, United States

**Keywords:** antimicrobial resistance, antibiotic stewardship, *Campylobacter*, surveillance, quinolones, tetracycline, aminoglycosides, macrolides, whole genome sequencing

## Abstract

*Campylobacter jejuni* and *Campylobacter coli* are important bacterial causes of human foodborne illness. Despite several years of reduced antibiotics usage in livestock production in the UK and US, high prevalence of antimicrobial resistance (AMR) persists in *Campylobacter*. Both countries have instigated genome sequencing-based surveillance programs for *Campylobacter*, and here we have identified AMR genes in 32,256 *C. jejuni* and 8,776 *C. coli* publicly available genome sequences to compare the prevalence and trends of AMR in *Campylobacter* isolated in the UK and US between 2001-2018. AMR markers were detected in 68% of *C. coli* and 53% of *C. jejuni*, with 15% of *C. coli* being multi-drug resistant (MDR) compared to only 2% of *C. jejuni*. The prevalence of aminoglycoside, macrolide, quinolone and tetracycline resistance remained fairly stable from 2001-2018 in both *C. jejuni* and *C. coli*, but statistically significant differences were observed between the UK and US. There was a statistically significant higher prevalence of aminoglycoside and tetracycline resistance for US *C. coli* and *C. jejuni*, and macrolide resistance for US *C. coli*. In contrast, UK *C. coli* and *C. jejuni* showed a significantly higher prevalence of quinolone resistance. Specific MLST clonal complexes (e.g. ST-353/464) showed >95% quinolone resistance. This large-scale comparison of AMR prevalence has shown that the prevalence of AMR remains stable for *Campylobacter* in the UK and the US. This suggests that antimicrobial stewardship and restricted antibiotic usage may help contain further expansion of AMR prevalence in *Campylobacter*, but are unlikely to reduce it in the short term.

## INTRODUCTION

Antibiotics are one of the great success stories of the 20^th^ century, changing medicine and agricultural practices (1). Unfortunately, the fairytale did not last, and over the last decades, antimicrobial resistance (AMR) has become a significant problem (2). The excessive and inappropriate use of antibiotics in human and veterinary medicine and their widespread use as growth promoters in agriculture has led to a rise of resistance to many classes of antibiotics, including last-resort, critically important antimicrobials for human medicine (3). This is now recognised as a serious and global crisis, especially given the lack of progress in developing novel antibiotics (4). The World Health Organisation declared AMR a severe threat to global public health in 2014 (5), and called upon the urgent introduction of multi-sectorial measures to prevent further development of AMR. This includes responsible use, which falls under antimicrobial stewardship (6, 7). One of the aspirations of reduced use of antimicrobials is that in the absence of antibiotic selection, susceptible isolates should have a fitness advantage over resistant isolates (8, 9), thereby restricting expansion of drug-resistant populations.

While phenotypic testing in the laboratory is still regarded as the gold standard, molecular tools such as those based on whole genome sequencing (WGS) are strong contenders to supersede phenotypic testing (10, 11), and have been successfully used in foodborne pathogens such as *Salmonella* and *Campylobacter* (12–15). However, harmonisation of databases and software tools is still required to avoid inter-laboratory reproducibility issues (16). Tools such as the NCBI AMRfinder software and curated Bacterial Antimicrobial Resistance Reference Gene Database at NCBI will certainly assist with such standardisation (17).

The bacterial pathogen *Campylobacter* is one of the leading causes of human bacterial diarrhoeal illness worldwide (18), and the European Food Safety Authority (EFSA) and the Centers for Disease Control (CDC) have reported similar levels of campylobacteriosis in the European Union and United States (19, 20). The causative agents, *C. jejuni* and *C. coli*, are commonly associated with poultry, wild birds, ruminants and pigs, with undercooked meat and cross-contamination seen as common causes of infection (21). The major categories of antibiotics used to treat *Campylobacter* are macrolides, such as erythromycin in humans. In contrast, aminoglycosides such as gentamicin and streptomycin, macrolides, quinolones such as ciprofloxacin and nalidixic acid, and tetracycline have been commonly used in agricultural and veterinary settings (22–24). The WHO has listed *Campylobacter* as a high-priority antibiotic-resistant pathogen due to the rapid rise in quinolone resistance in *Campylobacter* (2).

One of the requirements for unbiased monitoring of antimicrobial resistance levels in *Campylobacter* is the existence of large-scale surveillance studies with genome sequences and matching metadata deposited in public repositories such as NCBI and EMBL. Historical isolates and collections may suffer from biases introduced by a focus on isolates with ‘interesting’ phenotypes such as multiple antibiotic resistances. Only two countries have instigated large-scale genome sequencing-based surveillance of *Campylobacter:* the United Kingdom (UK) and United States of America (US). The UK surveillance has focused on clinical isolates (25, 26), whereas the US surveillance combines isolates from clinical, food and animal sources (27, 28).

In this study we have collected the available whole genome sequences from UK and US *C. jejuni* and *C. coli* isolates between 2001-2018, and have assessed the presence of genes conferring resistance to aminoglycosides, macrolides, quinolones and tetracyclines. We highlight the trends over time, differences between the two countries, and show the stability of the frequency of resistance to four antibiotic categories in *Campylobacter* in the UK and US which could raise concerns about future developments.

## RESULTS

### Comparison of genome assembly- and sequence read-based AMR marker detection

To assess the efficacy of the NCBI AMRfinder software tool with *Campylobacter* genome sequences, we compared it with a recent AMR analysis of 381 UK *C. jejuni* and *C. coli* isolates (13). Painset *et al*. (13) performed AMR marker detection directly on the Illumina sequencing reads using in-house scripts, whereas we used the NCBI AMRfinder software tool which requires genomes to be assembled first. There were no false-positives detected using the NCBI AMRfinder software for any of the antibiotic resistance classes (Table 1), and there was 100% agreement for detection of quinolone resistance (GyrA D87 mutation) and aminoglycoside resistance markers. Screening for tetracycline resistance resulted in 2/161 *tetO*-positive isolates that were initially reported negative for the *tetO* gene using NCBI AMRfinder. Further investigation of these two samples showed the *tetO* gene to be split over two contigs, which was detectable by secondary screening with BLAST using the Abricate software tool, and this was subsequently done as standard (Supplementary Table S1). Finally, macrolide resistance based on 23S rRNA gene mutations resulted in three negative samples previously reported positive (13). However, all three samples contained both the wild-type and mutated 23S rRNA alleles, suggesting only one or two of the three *C. jejuni* 23S rRNA genes mutated. Genome assembly algorithms are likely to ignore the minority gene variants, leading to these samples being tested negative. Finally, there were five genome assemblies that contained an *aph*(3’-IIIa aminoglycoside resistance gene (Table S1) not included in Painset *et al*. (13). Overall, there was very good concordance between the two tests, and we considered this a validation for testing *Campylobacter* genome assemblies with the NCBI AMRfinder software tool.

**Table 1.**
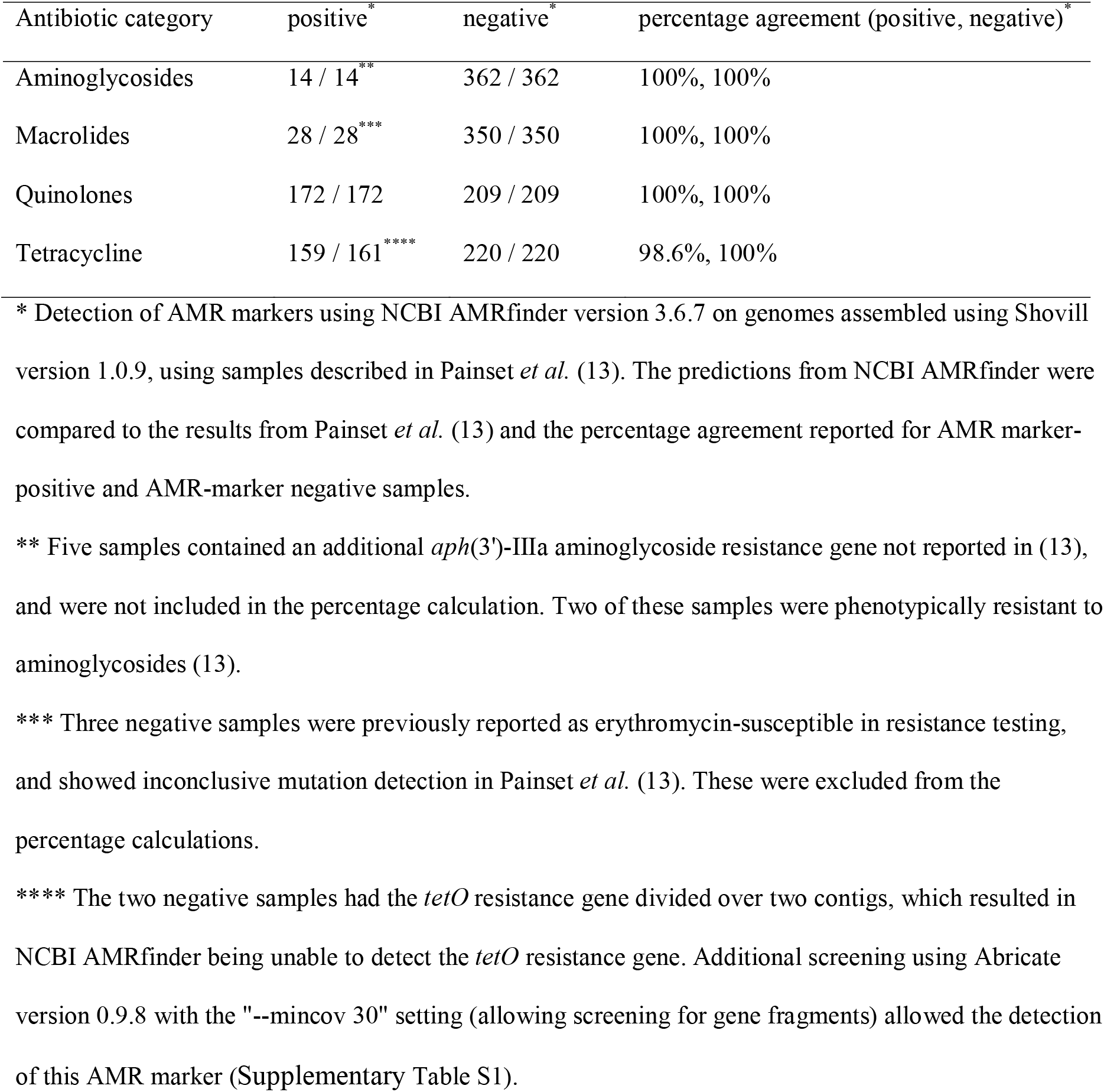
Validation of antimicrobial resistance marker detection in *Campylobacter* genomes using NCBI AMRfinder with assembled genomes, compared to the study of Painset *et al*. (13) which used detection of antimicrobial resistance markers with Illumina sequencing reads.

| Antibiotic category | positive<sup>*</sup> | negative<sup>*</sup> | percentage agreement (positive, negative)<sup>*</sup> |
| --- | --- | --- | --- |
| Aminoglycosides | 14 / 14<sup>**</sup> | 362 / 362 | 100%, 100% |
| Macrolides | 28 / 28<sup>***</sup> | 350 / 350 | 100%, 100% |
| Quinolones | 172 / 172 | 209 / 209 | 100%, 100% |
| Tetracycline | 159 / 161<sup>****</sup> | 220 / 220 | 98.6%, 100% |

\* Detection of AMR markers using NCBI AMRfinder version 3.6.7 on genomes assembled using Shovill version 1.0.9, using samples described in Painset *et al.* (13). The predictions from NCBI AMRfinder were compared to the results from Painset *et al.* (13) and the percentage agreement reported for AMR marker-positive and AMR-marker negative samples.
\*\* Five samples contained an additional *aph(3')-IIIa* aminoglycoside resistance gene not reported in (13), and were not included in the percentage calculation. Two of these samples were phenotypically resistant to aminoglycosides (13).
\*\*\* Three negative samples were previously reported as erythromycin-susceptible in resistance testing, and showed inconclusive mutation detection in Painset *et al.* (13). These were excluded from the percentage calculations.
\*\*\*\* The two negative samples had the *tetO* resistance gene divided over two contigs, which resulted in NCBI AMRfinder being unable to detect the *tetO* resistance gene. Additional screening using Abricate version 0.9.8 with the "--mincov 30" setting (allowing screening for gene fragments) allowed the detection of this AMR marker (Supplementary Table S1).

### Characteristics of a 2001-2018 *C. jejuni* and *C. coli* genome assembly database from public sources

A total of 32,256 *C. jejuni* and 8,776 *C. coli* genomes were included in our study. The distribution over the year categories is shown in Figure 1A; for *C. coli*, approximately 400 UK samples were added for each of the 2015-2018 years, whereas the number of US *C. coli* samples increased from approximately 500 samples in 2015 to just over 2,000 in 2018. Similar trends were observed for *C. jejuni*, where the number of UK samples varies between approximately 2,000 and 4,000 per year, while the number of US samples increased from approximately 1,000 in 2015 to just under 4,000 in 2018. The samples from 2001-2014 were combined, as these did not have consistent availability of samples, with yearly numbers varying between 0-350 (*C. coli*, UK), 7-704 (*C. coli*, US), 10-1,881 (*C. jejuni*, UK) and 0-277 (*C. jejuni*, US) (Supplementary Figures S1, S2).

**Figure 1.**
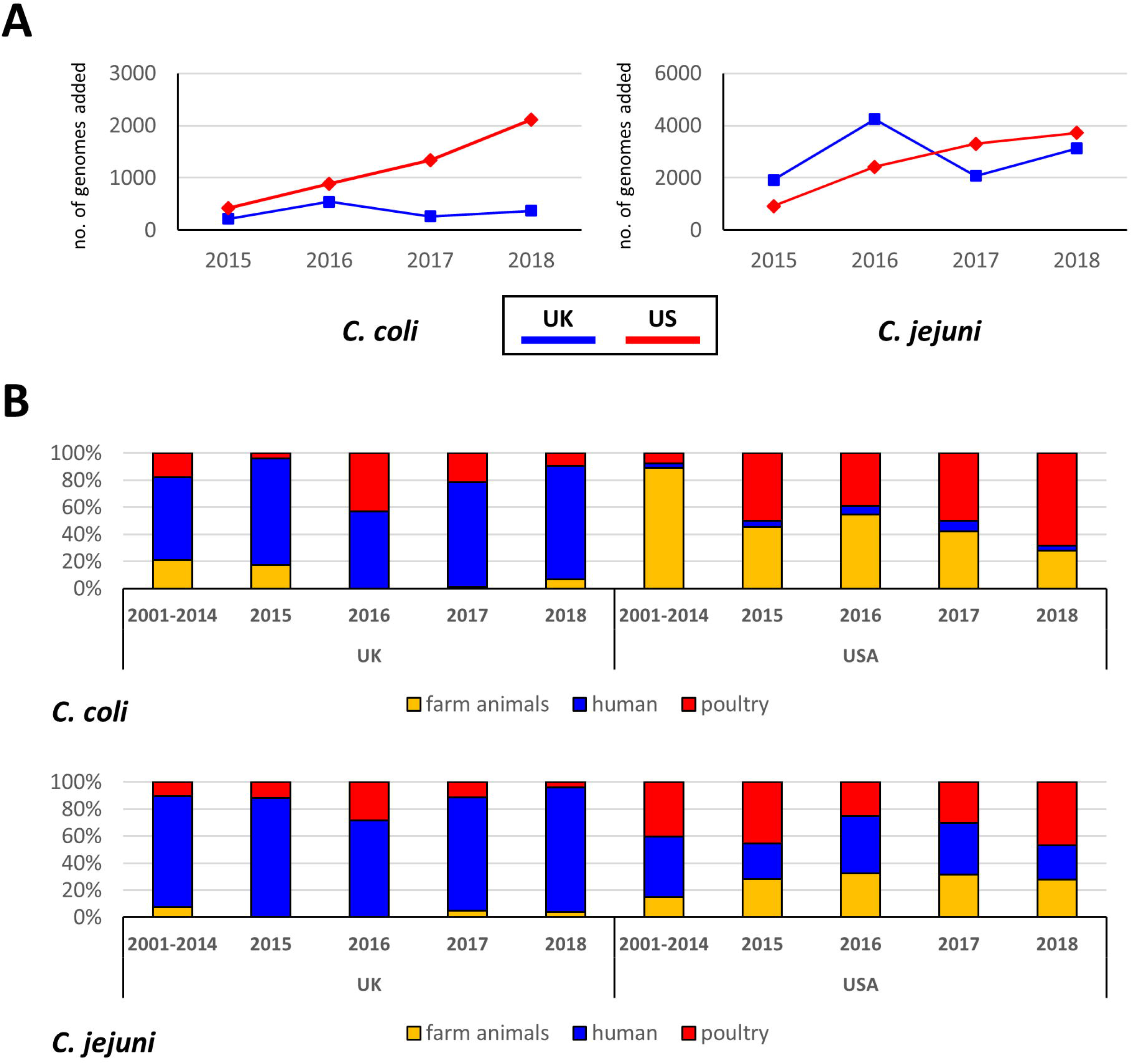
Breakdown of the UK and US *C. coli* and *C. jejuni* samples obtained in surveillance programs from 2015-2018. A) Number of genome sequences per individual year from 2015-2018. The number of US genome sequences shows steady growth each year, whereas the number of UK genome sequences varies per year without a clear trend. B) Comparison of the isolation source category of the UK and US samples from 2015-2018, with the 2001-2014 historical isolates included. Within the UK samples, the human source category is dominant, consistent with the major surveillance category being the Oxfordshire sentinel surveillance (25, 26), while US isolates have a much higher contribution of farm animals (pigs, ruminants) and poultry (27, 28).

The *C. jejuni* and *C. coli* samples were split into several groups: UK and US-derived samples, and within each historical samples from 2001-2014 versus the 2015-2018 individual years, and the three source categories (farm animals, human, poultry). Comparison of these categories (Figure 1B) showed that UK samples were dominated by human isolates (approximately 70% for *C. coli* vs 5% for *C. coli* from the US), with farm animal samples poorly represented. In contrast, US samples showed a better representation of farm animal samples for both *C. jejuni* and *C. coli*, especially in samples derived from 2001-2014.

### Distribution of antimicrobial resistance markers in *C. jejuni* and *C. coli*

The antimicrobial resistance profiles were determined for all the *C. jejuni* and *C. coli* genomes using NCBI AMRfinder. The presence of any of the 44 resistance markers was translated into aminoglycoside resistance (20 markers), macrolide resistance (9 markers), quinolone resistance (10 markers) or tetracycline resistance (5 markers). Table 2 shows the breakdown of the number of isolates predicted to be resistant for both *C. coli* and *C. jejuni*.

**Table 2.**
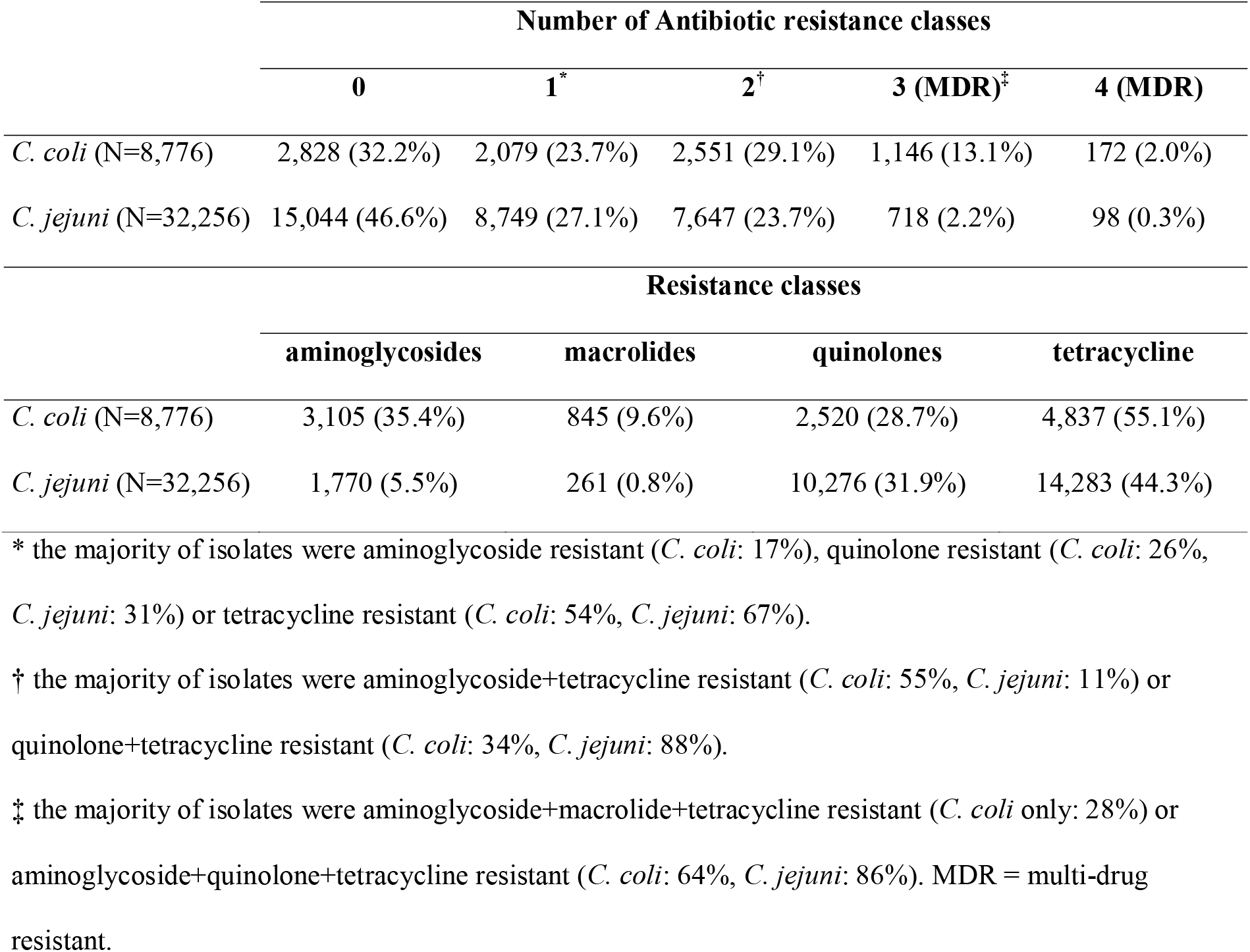
Distribution of antibiotic resistance classes in *C. coli* and *C. jejuni* genome sequences (2001-2018 combined).

In *C. coli*, 15.1% of samples were predicted to be multi-drug resistant (MDR, resistant to three or more classes of antibiotics), whereas only 2.5% of *C. jejuni* were predicted to be MDR. Similarly, almost half of *C. jejuni* samples (46.6%) did not have any resistance marker compared to one-third of *C. coli* samples (32.2%). Approximately half of the *C. coli* and *C. jejuni* isolates were predicted to be resistant to tetracycline (55.1% and 44.3%, respectively), while about one-third of *C. coli* and *C. jejuni* were predicted to be resistant to quinolones (28.7% and 31.9%, respectively). Notable differences were seen in aminoglycoside resistance (*C. coli* 35.4% vs *C. jejuni* 5.5%) and macrolide resistance (*C. coli* 9.6% vs *C. jejuni* 0.8%) (Table 2). Of note, the prevalence of gentamicin resistance was 213/8,776 (2.4%) for *C. coli* and 127/32,256 (0.4%) for *C. jejuni*, with most gentamicin resistance markers detected in US samples (Supplementary Table S1).

### Comparison of AMR levels between the UK and US (2001-2018)

The UK and US are currently the two countries which do large-scale genome sequencing-based surveillance of *Campylobacter* (14, 26), and this allows for insight in the dynamics of antibiotic resistance in these two industrialised countries over time. The years covering 2015-2018 were used individually while combining the 2001-2014 samples to function as a possible baseline for comparative purposes. The predicted resistance to aminoglycosides, macrolides, quinolones and tetracycline were plotted out for the UK and US isolates and compared for trends (Figure 2), with the individual years from 2001-2018 shown in Supplementary Figure S1 and S2.

**Figure 2.**
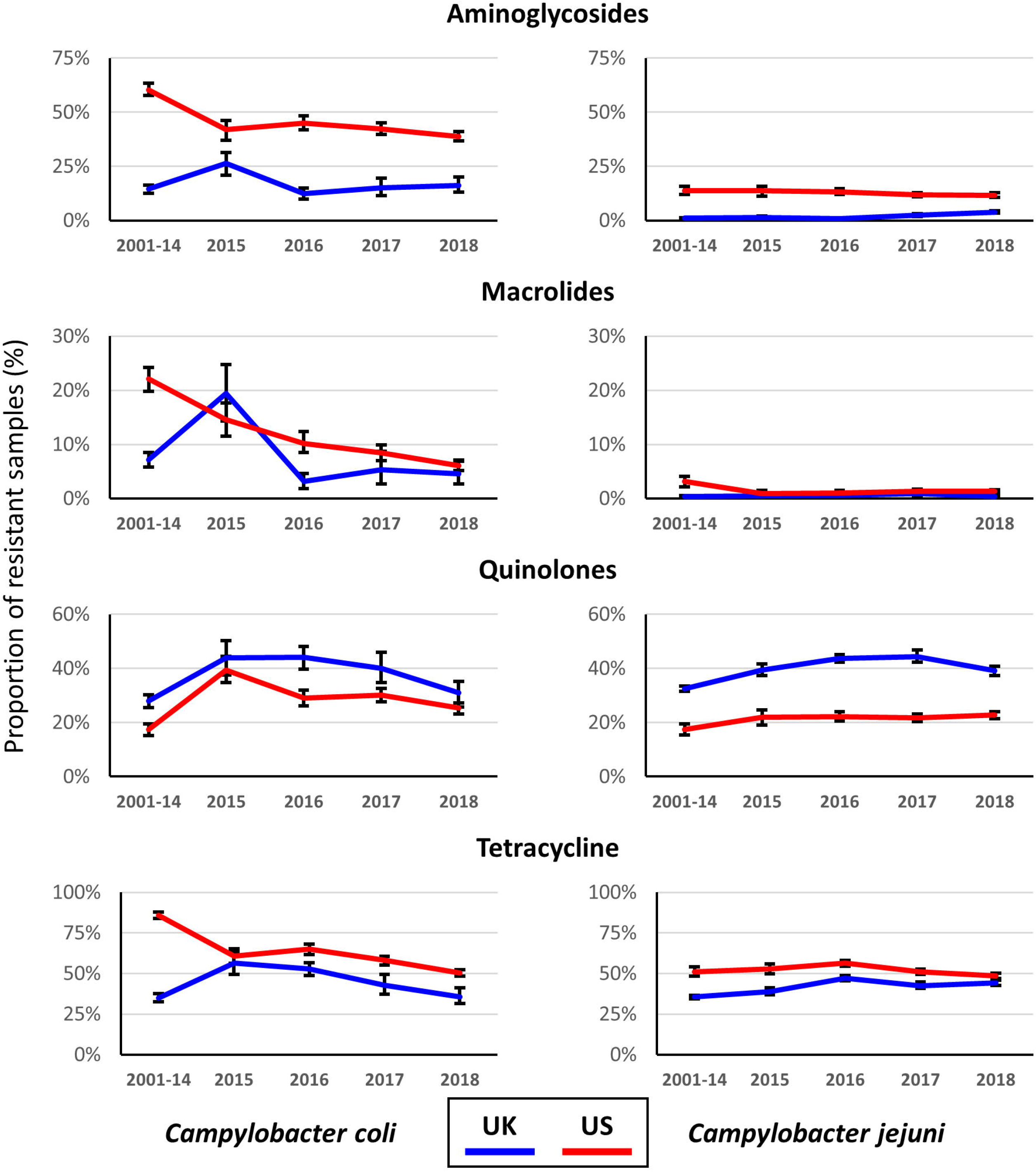
Comparison of the proportion of UK and US *C. coli* and *C. jejuni* samples resistant to aminoglycosides, macrolides, quinolones and tetracycline for individual years from 2015-2018, with the average of the 2001-2014 period provided for comparative purposes. Error bars show the 2.5% and 97.5% quantiles, based on 500 bootstraps.

The general trend for both *C. coli* and *C. jejuni* was that the proportion of aminoglycoside and tetracycline-resistant isolates was significantly higher in the US than in the UK between 2015-2018, while the prevalence of quinolone resistance was significantly lower in the US (Figure 2), and prevalence of macrolide resistance was similar for both countries for *C. jejuni*, and was higher for US *C. coli* for three of the four years, but lower in 2015 (Figure 2). Approximately 40% of the US *C. coli* isolates were predicted to be resistant to aminoglycosides (range 39-45%), whereas in the UK *C. coli* isolates this was approximately 20% (range 12-26%). For *C. jejuni*, the prevalence of aminoglycoside resistance was between 12-14% for the US isolates compared to 1-4% for the UK isolates. Differences were less pronounced for tetracycline resistance for *C. jejuni* (range 39-46% for UK vs 48-56% for the US samples), while in *C. coli* this varied between 36-56% for the UK isolates, and 50-65% for the US isolates. Macrolide resistance was present in much higher levels in *C. coli* in both the UK and the US isolates, ranging from 3-19% in the UK isolates and 6-14% in the US *C. coli* isolates, while it was between 0-1% in the UK and the US *C. jejuni* isolates. Finally, the trend was reversed for quinolone resistance, with a higher proportion of the UK isolates predicted to be quinolone-resistant. For *C. coli*, the proportion of quinolone-resistant UK isolates was 31-44% for UK isolates versus 25-39% for US isolates, while for *C. jejuni* the proportion was 39-44% for UK isolates and 22-23% for US isolates. For each of the *Campylobacter* species and antibiotic class, statistically significant differences were generally observed between countries and year (Supplementary Table S2).

Next to the differences between the two countries, we also observed that the only antibiotic for which there was a clear decrease in resistance from 2015 to 2018 was macrolide resistance (Figure 2), as it decreased from 15% to 6% in the US isolates, and from 19% to 5% in the UK isolates. For the other three antibiotic classes, there was little change from 2015-2018 and especially in *C. coli*, the proportion of resistant isolates remains high for aminoglycoside resistance (US), quinolone resistance (UK, US) and tetracycline resistance (UK, US). For *C. jejuni*, the levels of quinolone resistance are especially of concern in the UK.

### Associations between isolation source and MLST genotypes and AMR markers in *C. coli* and *C. jejuni*

Due to the pronounced differences in isolation source between the UK and US (Figure 1B), we investigated whether there were differential contributions to the proportions of antibiotic-resistant isolates for *C. coli* and *C. jejuni* isolates from 2015-2018 (Figure 3A). There was no clear pattern of overrepresentation of any isolation source for the AMR isolates in *C. coli*., However, the low number of farm animal isolates from the UK may hide some effects. Similarly, for *C. jejuni* there was no clear link with any of the three source categories (Figure 3A).

**Figure 3.**
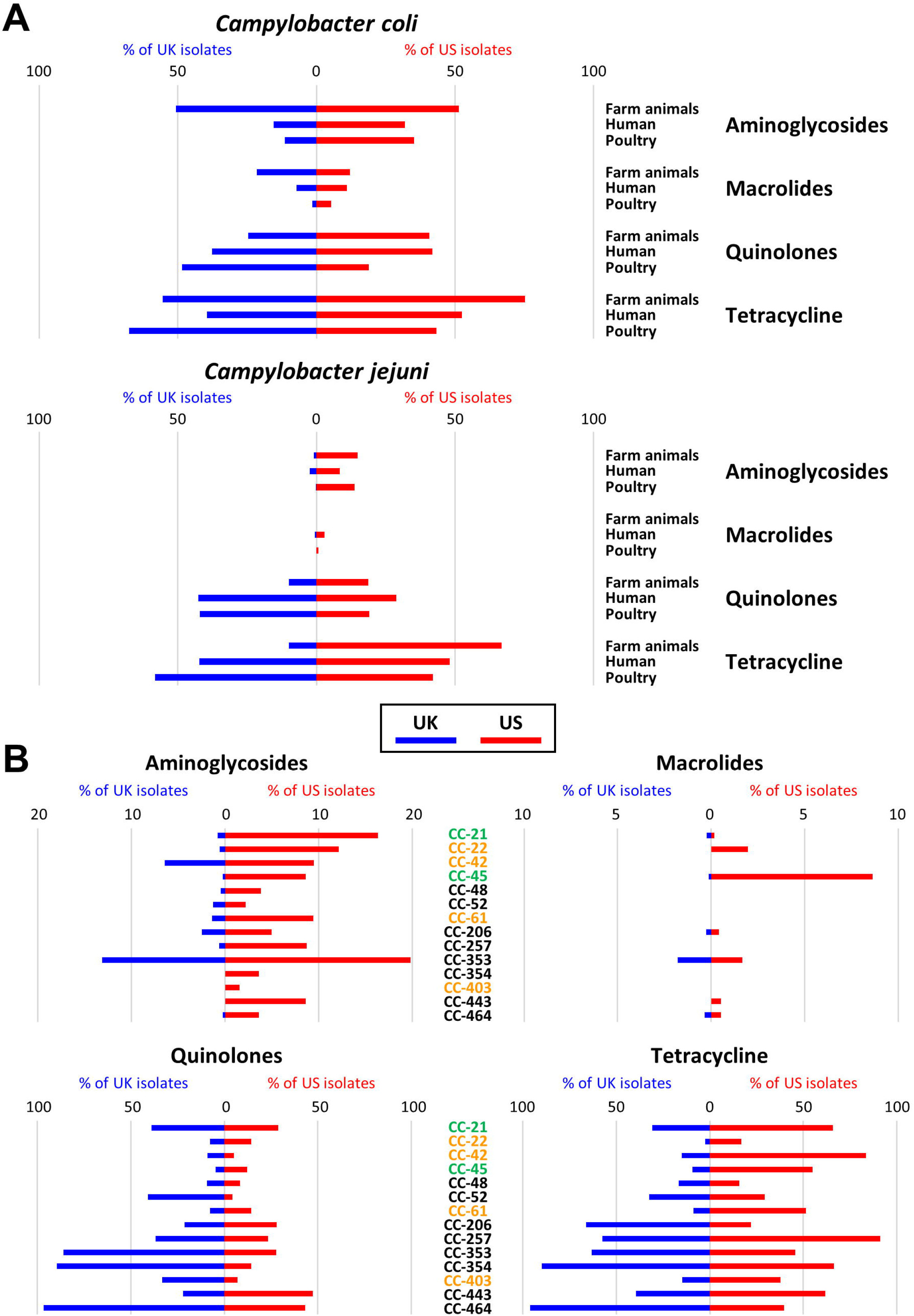
Tornado plots displaying the relative contributions of the source categories (A) and *C. jejuni* MLST clonal complexes (B) to the individual antibiotic classes (aminoglycosides, macrolides, quinolones and tetracycline). The horizontal axis shows the percentage UK samples from 100 to 0 on the left half, and the percentage US samples on from 0 to 100 on the right half for each of the categories. Only the samples form the years 2015-2018 have been included for this figure.

We also investigated whether specific multilocus sequence types (MLST) clonal complexes (CC) were associated with specific antibiotic resistances in the UK and US isolates. For *C. coli*, 97.3% of the isolates are from CC-828. For *C. jejuni*, we looked at the clonal complexes for which both for the UK and US there were at least >100 samples available for 2015-2018 (Figure 3B). For aminoglycoside resistance, there was a spread over all the major clonal complexes for US isolates, with CC-21 and CC-353 having the highest proportion of resistant isolates (17-20%). In contrast, for the UK isolates, aminoglycoside resistance was primarily found in CC-353 and CC-42. The CC-353 clonal complex was also involved in macrolide resistance, although this plays a minor role in the *C. jejuni* isolates, with CC-45 showing the highest proportion of resistant isolates in the US. For both quinolone resistance and tetracycline resistance, all major MLST genotypes contributed to resistance, but the major contribution to quinolone resistance in UK isolates came from CC-353, CC-354 and CC-464, where almost all isolates were predicted to be quinolone resistant, consistent with earlier reports (26, 29). Finally, the highest proportions of tetracycline resistance were associated with CC-354 and CC-464 for UK isolates, and CC-257 and CC-42 for US isolates. Most of the clonal complexes reported here are primarily associated with poultry and human infections, except for CC-42 associated with cattle.

## DISCUSSION

In this study we have exploited the possibilities afforded by the active genome sequencing-based surveillance for *Campylobacter* in the UK and US, to determine the trends in AMR over recent years, and compare these trends within and between the two countries. Similar studies have previously been conducted in individual countries, but often at a smaller scale than employed in this study, or focusing on a shorter time period. For instance, a recent UK study investigated 528 human isolates from 2015-2016 (13), while a 2018 US-based study focused on 589 isolates from 2015 (14). These, and other genome sequencing-based studies have all shown that identifying resistance markers in genome sequences matches well with phenotypic resistance. Hence we are confident that our study appropriately represents AMR trends detected in 32,256 *C. jejuni* and 8,876 *C. coli* samples, as listed in Supplementary Table S1.

Monitoring of AMR on this scale requires active surveillance with sequencing data and metadata being released in the public domain. We have combined the information present in the Genbank database, which contains both US and UK samples, and the Campylobacter PubMLST database which includes a large number of UK isolates. Our dataset is based on a de-duplicated, quality controlled set of genome sequences with strict requirements for available metadata. The criteria for isolate source, year and country data allowed direct comparison between the UK and US *Campylobacter* samples for *C. coli* and *C. jejuni* (Figure 1B). The UK dataset leans heavily on the Oxfordshire Sentinel Surveillance (25, 26), which explains the dominance of human samples in the UK dataset in each individual year. This contrasts with the US dataset, which has an equal distribution over the three source categories. While this is not ideal for comparative purposes, we do note that for *C. jejuni* the distribution over the dominant MLST clonal complexes was very similar (Figure 3), with the major differences being that CC-257 and CC-464 were prevalent in a higher proportion in UK samples (5% and 6% more of the total number of samples). In comparison, CC-353 was more prevalent in US samples (7% more). All three clonal complexes are primarily associated with poultry, and hence for *C. jejuni*, we concluded that the datasets can still be compared. For *C. coli* this was more difficult to assess, as most samples cluster in the agricultural Clade 1A (30, 31) which only contains CC-828 and a minority of samples not part of this clonal complex. The individual MLST sequence types (STs) may primarily represent the STs prevalent in the two countries, and may be affected by the differences in isolation sources between the samples from the two countries (Fig. 1B).

In this study we have focused on four classes of antibiotics, due to their relevance in *Campylobacter* and their usage in the categories of hosts investigated. The macrolide erythromycin and the fluoroquinolone ciprofloxacin are used to treat human infections with *Campylobacter* (22). In contrast, all four classes of antibiotics are used therapeutically in the veterinary sector for livestock, albeit not for control of *Campylobacter*, as the usage is associated with other infectious diseases in livestock (32). As highlighted earlier, the WHO has listed *Campylobacter* as a high-priority antibiotic-resistant pathogen (2), especially linked to fluoroquinolone resistance in *Campylobacter*. Antimicrobial resistance in *Campylobacter* spp. is a combination of *de novo* point mutations associated with quinolone and macrolide resistance, and genetic exchange on plasmids and other mobile elements for aminoglycoside and tetracycline resistance genes (33). In addition, the natural competence of *C. jejuni* and *C. coli* (34, 35) may allow further dissemination of resistance through populations. Although the usage of antibiotics has been limited in the UK and the US for a considerable period of time (32, 36, 37), there is no apparent reduction visible in the prevalence of resistance in the large numbers of *C. jejuni* and *C. coli* samples investigated here (Figure 2). Our statistical analyses suggest that there may be small increases in *C. jejuni* resistance to some antibiotics in recent years, when compared compared the levels of 2014 and earlier (Supplementary Table S2). However, due to the composition of the dataset and its retrospective nature, this is something that requires a prospective study to further investigate, which falls outside of the aims and scope of the work presented here. A possible hypothesis for the observations could be that resistance to especially tetracyclines, aminoglycosides and quinolones is not detrimental to *Campylobacter*, or not sufficiently detrimental to allow out-competition by susceptible isolates. Conversely, resistance may even be associated with advantageous phenotypes, as the gyrase mutations associated with quinolone resistance gave rise to increases in virulence phenotypes in *C. jejuni* (38, 39), while *tetO*-containing plasmids are ubiquitous and stable in *C. jejuni* and *C. coli* (40, 41). Aminoglycoside resistance genes are especially widespread in *C. coli*, and their association with mobile elements (33, 42) may assist their stability in the populations.

The similarities between the UK and US datasets suggest that the situation in other industrialised countries may be similar, but this will require more intensive surveillance programs and will need to balance clinical, food and agricultural samples, including sewage and water sources. A recent study reported that fluoroquinolone usage in pigs and poultry in France from 2011-2018 decreased by >70% in poultry, and almost 90% in pigs. However, resistance to ciprofloxacin increased in poultry *Campylobacter* isolates from 50% to approximately 60%, whereas in *Escherichia coli* prevalence of ciprofloxacin resistance was stable in broiler and pig isolates, but decreased in turkey isolates (43). Similar studies have been done in other countries, but often are based on small or limited datasets where selection bias may affect the results.

The lack of reduction of the proportion of AMR isolates in *Campylobacter* from both countries does suggest that reduced usage of antimicrobials, such as mandated by antimicrobial stewardship, will not be sufficient to reduce the incidence of AMR in *Campylobacter*, although it may well contribute positively to reigning in further increases of AMR. For clinical purposes, macrolides such as erythromycin are still useful for treating *C. jejuni* infections, but their efficacy may be reduced for *C. coli*. In contrast, the high level of quinolone resistance in both *C. jejuni* and *C. coli* does not bode well for the future efficacy of ciprofloxacin treatment of *Campylobacter* infections in humans. With regard to aminoglycosides and tetracyclines, the latter is not recommended due to the high level of resistance, whereas with aminoglycosides gentamicin can still be used for now, as prevalence of gentamicin resistance is still low.

Taken together, the data presented here strongly suggest that reduced usage of antibiotics has not resulted in a significant reduction of antimicrobial resistance in *Campylobacter*, which is of considerable public health and economic concern. Changes in agricultural practices, slaughter and retail will need to be substantial to reduce the overall prevalence of *Campylobacter*. These efforts should lessen the need for antibiotic usage to achieve the goals of antimicrobial stewardship. Similar comparative studies could be done in other countries and other foodborne zoonotic infections to assess whether this situation is unique for *Campylobacter* or mirrored in other pathogenic bacteria.

## MATERIALS AND METHODS

### *Campylobacter* genome assemblies and metadata categories included in this study

A total of 44,751 *C. jejuni* and 12,709 *C. coli* genome assemblies were collected from the Genbank and Campylobacter PubMLST databases, and were coupled to metadata from the Genbank files and from PubMLST. Genome assemblies were obtained from the NCBI database using ncbi-genome-download version 0.2.11 (https://github.com/kblin/ncbi-genome-download/), and supplemented with genome sequences from the *Campylobacter* pubMLST website (http://pubmlst.org/campylobacter/) (44). Metadata were extracted from Genbank flat files and the NCBI Pathogens database (https://www.ncbi.nlm.nih.gov/pathogens). All genome assemblies were screened for assembly statistics using Quast version 4.5 (45), and genome assemblies were excluded if failing two or more of the following criteria: number of contigs ≤ 200, N50 ≥ 25 kb, L50 ≤ 25 contigs, largest contig ≥ 50 kb or the number of Ns per 100 kb ≥ 50. Genomes with a total size outside 1.4 Mbp and 2.1 Mbp were automatically excluded. Duplicate entries were removed by comparing sample names, assembly statistics such as N50, L50 and genome size and metadata such as year, source and country. After deduplication, samples were subsequently only included if the following metadata were available: isolation source, year of isolation, and isolated in the UK or US. Isolation sources were combined to give three main categories: farm animals (pigs, cattle, sheep, goat, including milk and meat samples), human (clinical isolates) and poultry (chicken, turkey, including meat samples). Samples representing environmental, farm, generic “food” and wild birds were excluded, as were samples lacking other metadata. This resulted in the exclusion of 6,327 *C. jejuni* and 877 *C. coli* samples with missing information on country, year or isolation source; 1,634 *C. jejuni* and 142 *C. coli* were not from the UK or US; 3,116 *C. jejuni* and 2,086 *C. coli* were from outside the 2001-2018 period studied here, and 1,418 *C. jejuni* and 828 *C. coli* were not from animal, ruminant, poultry or human sources. We focused on 2015-2018 as individual years as both UK and US doing surveillance projects, while historical samples were categorised as 2001-2014. Overall, we analysed a total of 41,032 genome sequences, represented by 32,256 *C. jejuni* and 8,776 *C. coli* genomes (Supplementary Table S1).

### Screening for antimicrobial resistance markers

Genome assemblies were screened for AMR markers using the NCBI AMRfinder software tool version 3.1.1b (17), with the October 2019 database, and the nucleotide setting (-n) and the organism switch (-O Campylobacter) which includes screening for point mutation-based resistances. Genome assemblies were also screened using Abricate version 0.9.8 (https://github.com/tseemann/abricate/) and the NCBI database, which screens for AMR genes, but not point mutation-based resistances. Resistance genes were categorised into four types of resistance: aminoglycoside resistance [*aph*(2”)-Ia, *aph*(2”)-If, *aph*(2”)-If2, *aph*(2”)-Ig, *aph*(2”)-Ih, *aph*(2”)-IIa, *aph*(2”)-IIIa, *aph*(2”)-IVa, *apmA*, *aac*(6’)-Ie, *aac*(6’)-Im, *aad9*, *aadE*, *aadE-Cc*, *ant*(6)-Ia, *aph*(3’)-IIa, *aph*(3”)-Ib, *aph*(3’)-IIIa, *aph*(3’)-VIIa, spw], macrolide resistance [23S_A2074C, 23S_A2074G, 23S_A2074T, 23S_A2075G, *cfr*(C), *erm*(36), *erm*(B), *erm*(C), *erm*(F)], quinolone resistance [GyrA_D90N, GyrA_D90Y, GyrA_P104S, GyrA_T86A, GyrA_T86I, GyrA_T86K, GyrA_T86V, *qnrB*, *qnrD*, *qnrD1*] and tetracycline resistance [*tet*(32), *tet*(L), *tet*(O), *tet*(W), *tet*(X)]. Samples were considered multi-drug resistant (MDR) when containing resistance markers for three or more classes of antibiotics.

### Comparison of genome assembly-based screening for AMR genes with sequencing-read based screening

The Supplementary Data presented in Painset *et al*. (13) contained 381 accession numbers to FASTQ read files of UK *Campylobacter* samples used for screening for antimicrobial resistance genes. The FASTQ files were downloaded from the Sequence Read Archive using fastq-dump from the SRA toolkit (https://github.com/ncbi/sra-tools), and genomes assembled using Spades version 3.14 (46) via the Shovill version 1.0.9 tool and standard settings (https://github.com/tseemann/shovill/). All genomes passed the Quast QC, and were screened using the NCBI Amrfinder tool version 3.6.7 (17) with the nucleotide setting (“-n”) and the organism switch (“-O Campylobacter”) as described above. All genome assemblies were also screened using Abricate version 0.9.8 (https://github.com/tseemann/abricate/) and the NCBI database with a minimum coverage of 30% (--mincov 30) to check for the *tetO* tetracycline resistance gene being split over two or more contigs. These 381 genome assemblies have not been included in the other analyses presented here.

### Statistical analysis of association between resistance markers and descriptor variables (country, year and isolation source)

The presence or absence of resistance for each of the four classes of antibiotics was assessed using a generalized linear model with a logit link function (binomial family). Country, year and isolation source of each sample were considered as explanatory variables. Interactions between predictors were not considered. Bootstrapping with 500 repeats was carried out to estimate the 95% confidence interval (CI) of the prevalence of resistance for each country and year. All statistical analyses were carried out in R (version 4.0.3) (47), and the data are presented in Supplementary Table S2.

### Data description

All genome sequences used in this study are available from the Genbank/EMBL/DDBJ databases or the Campylobacter PubMLST website (https://pubmlst.org/organisms/campylobacter-jejunicoli/). The assembly accession numbers (NCBI Genome) or genome ID numbers (Campylobacter PubMLST) are listed in Supplementary Table S1, together with the metadata used and the AMR gene data.

## ACKNOWLEDGMENTS

We gratefully acknowledge the efforts of the surveillance programs in the US and UK, and their continued contribution to data sharing by making the genome sequences and partial metadata publicly available. Part of this work was supported by funding from the European Union’s Horizon 2020 Research and Innovation programme under grant agreement No 773830: One Health European Joint Programme (WorldCOM). The US whole genome sequencing work was supported by the National Institutes of Health/Food and Drug Administration under award number 5U19FD007113-02, whereas UK *Campylobacter* surveillance was supported by the UK Food Standards Agency. This publication made use of the PubMLST website (http://pubmlst.org/) developed by Keith Jolley and sited at the University of Oxford. The development of that website was funded by the Wellcome Trust.

## SUPPLEMENTARY INFORMATION

**Supplementary Table S1.** Overview of *C. jejuni* and *C. coli* genomes used in this study, with the Genbank or Campylobacter PubMLST accession numbers, metadata (source, MLST clonal complex, MLST sequence type), whether they are resistant to aminoglycosides, macrolides, quinolones or tetracycline, and which individual resistance marker they are positive for.

**Supplementary Table S2.** Results from the generalized linear regression with a logit link function of AMR prevalence for each of the four antibiotic classes (aminoglycosides, macrolides, quinolones and tetracycline) for *C. coli* and *C. jejuni* isolates. Country, source and year categories are considered as fixed effects; UK, human source and the pre-2015 categories are used as reference. Odds can be calculated from the exponential of the regression coefficients shown. Statistically significant increases from the reference are highlighted in red, decreases in green.

**Supplementary Figure S1.** Comparison of the proportion of UK and US *C. coli* samples resistant to aminoglycosides, macrolides, quinolones and tetracycline for individual years from 2001-2018.

**Supplementary Figure S2.** Comparison of the proportion of UK and US *C. jejuni* samples resistant to aminoglycosides, macrolides, quinolones and tetracycline for individual years from 2001-2018.

