## Supplementary Table S2, Figure S1, Figure S2 for "Genomic screening of antimicrobial resistance markers in UK and US *Campylobacter* isolates highlights stability of resistance over an 18 year period"

Running title: Stability *Campylobacter* AMR over 18 yr period

Keywords: antimicrobial resistance, antibiotic stewardship, Campylobacter, surveillance, quinolones, tetracycline, aminoglycosides, macrolides, whole genome sequencing

**Supplementary Table S1.** Overview of *C. jejuni* and *C. coli* genomes used in this study, with the Genbank or Campylobacter PubMLST accession numbers, metadata (source, MLST clonal complex, MLST sequence type), whether they are resistant to aminoglycosides, macrolides, quinolones or tetracycline, and which individual resistance marker they are positive for [Excel File].

**Table S2.** Results from the generalized linear regression with a logit link function of AMR prevalence for each of the four antibiotic classes (aminoglycosides, macrolides, quinolones and tetracycline) for *C. coli* and *C. jejuni* isolates. Country, source and year categories are considered as fixed effects; UK, human source and the pre-2015 categories are used as reference. Odds can be calculated from the exponential of the regression coefficients shown. Statistically significant increases from the reference are highlighted in red, decreases in green.

| Explanatory variable | | Regression coefficient | Standard error | Z value | Pr(>\|z\|) |
|---|---|---|---|---|---|
| ***Campylobacter coli / Aminoglycosides*** | | | | | |
| Intercept | | -1,822081 | 0,069152 | -26,348895 | < 2e-16 |
| Country | UK | 0 | | | |
| | USA | 1,249063 | 0,079098 | 15,79127 | < 2e-16 |
| Source | Human | 0 | | | |
| | Poultry | 0,185077 | 0,086271 | 2,145302 | 0,031929 |
| | Farm animal | 0,922235 | 0,087311 | 10,562622 | < 2e-16 |
| Year | Pre-2015 | 0 | | | |
| | 2015 | -0,056304 | 0,101024 | -0,557333 | 0,5773 |
| | 2016 | -0,209148 | 0,078598 | -2,660973 | 0,007792 |
| | 2017 | -0,203846 | 0,076565 | -2,662407 | 0,007758 |
| | 2018 | -0,241704 | 0,072332 | -3,341586 | 0,000833 |
| ***Campylobacter coli / Macrolide*** | | | | | |
| Intercept | | -2,306224 | 0,090055 | -25,609075 | < 2e-16 |
| Country | UK | 0 | | | |
| | USA | 0,884907 | 0,123122 | 7,187251 | 6.61e-13 |
| Source | Human | 0 | | | |
| | Poultry | -0,661522 | 0,136626 | -4,841857 | 1.29e-06 |
| | Farm animal | 0,097725 | 0,128726 | 0,759175 | 0,447748 |
| Year | Pre-2015 | 0 | | | |
| | 2015 | 0,212842 | 0,125891 | 1,69068 | 0,090898 |
| | 2016 | -0,655531 | 0,119539 | -5,483813 | 4.16e-08 |
| | 2017 | -0,694965 | 0,115853 | -5,998694 | 1.99e-09 |
| | 2018 | -0,917685 | 0,113998 | -8,050017 | 8.28e-16 |
| ***Campylobacter coli / Quinolone*** | | | | | |
| Intercept | | -0,916714 | 0,057221 | -16,020549 | < 2e-16 |
| Country | UK | 0 | | | |
| | USA | -0,294318 | 0,073922 | -3,981448 | 6.85e-05 |
| Source | Human | 0 | | | |
| | Poultry | -0,690244 | 0,077311 | -8,92809 | < 2e-16 |
| | Farm animal | -0,159259 | 0,080711 | -1,973194 | 0.05 |
| Year | Pre-2015 | 0 | | | |
| | 2015 | 1,0272 | 0,096029 | 10,69682 | < 2e-16 |
| | 2016 | 0,783259 | 0,075276 | 10,405171 | < 2e-16 |
| | 2017 | 0,740323 | 0,077148 | 9,596095 | < 2e-16 |
| | 2018 | 0,547365 | 0,073438 | 7,453435 | 9.09e-14 |
| ***Campylobacter coli / Tetracycline*** | | | | | |
| Intercept | | -0,521043 | 0,054118 | -9,627851 | < 2e-16 |
| Country | UK | 0 | | | |

| | | | | | |
|---|---|---|---|---|---|
| | USA | 0,613287 | 0,069058 | 8,880791 | < 2e-16 |
| Source | Human | 0 | | | |
| | Poultry | 0,078275 | 0,071239 | 1,098775 | 0.27187 |
| | Farm animal | 1,159653 | 0,07715 | 15,031137 | < 2e-16 |
| Year | Pre-2015 | 0 | | | |
| | 2015 | 0,100718 | 0,09698 | 1,038542 | 0.29902 |
| | 2016 | 0,19793 | 0,073893 | 2,678589 | 0.00739 |
| | 2017 | -0,188263 | 0,074355 | -2,531961 | 0.01134 |
| | 2018 | -0,405295 | 0,069232 | -5,854121 | 4.8e-09 |
| ***Campylobacter jejuni / Aminoglycosides*** | | | | | |
| Intercept | | -4,285236 | 0,072056 | -59,471228 | < 2e-16 |
| Country | UK | 0 | | | |
| | USA | 1,979807 | 0,072473 | 27,317782 | < 2e-16 |
| Source | Human | 0 | | | |
| | Poultry | 0,18934 | 0,063193 | 2,99623 | 0.00273 |
| | Farm animal | 0,334018 | 0,067496 | 4,94873 | 7.47e-07 |
| Year | Pre-2015 | 0 | | | |
| | 2015 | 0,210609 | 0,109198 | 1,928694 | 0.05377 |
| | 2016 | 0,151285 | 0,087529 | 1,728399 | 0.08392 |
| | 2017 | 0,175373 | 0,08712 | 2,013003 | 0.04411 |
| | 2018 | 0,253927 | 0,08369 | 3,034126 | 0.00241 |
| ***Campylobacter jejuni / Macrolide*** | | | | | |
| Intercept | | -5,076742 | 0,126506 | -40,130421 | < 2e-16 |
| Country | UK | 0 | | | |
| | USA | 1,8675 | 0,144005 | 12,968332 | < 2e-16 |
| Source | Human | 0 | | | |
| | Poultry | -1,583257 | 0,19354 | -8,180532 | 2.83e-16 |
| | Farm animal | -16,951664 | 267,949287 | -0,063264 | 0.950 |
| Year | Pre-2015 | 0 | | | |
| | 2015 | -0,332244 | 0,272584 | -1,218867 | 0.223 |
| | 2016 | -0,271909 | 0,19116 | -1,422418 | 0.155 |
| | 2017 | -0,083558 | 0,187026 | -0,446774 | 0.655 |
| | 2018 | -0,188755 | 0,187048 | -1,009126 | 0.313 |
| ***Campylobacter jejuni / Quinolone*** | | | | | |
| Intercept | | -0,644846 | 0,0219 | -29,444562 | < 2e-16 |
| Country | UK | 0 | | | |
| | USA | -0,62379 | 0,031974 | -19,509347 | < 2e-16 |
| Source | Human | 0 | | | |
| | Poultry | -0,386911 | 0,033317 | -11,612996 | < 2e-16 |
| | Farm animal | -0,907609 | 0,048464 | -18,727567 | < 2e-16 |
| Year | Pre-2015 | 0 | | | |
| | 2015 | 0,291209 | 0,046193 | 6,304132 | 2.9e-10 |
| | 2016 | 0,458931 | 0,034489 | 13,306446 | < 2e-16 |
| | 2017 | 0,423712 | 0,039518 | 10,722006 | < 2e-16 |
| | 2018 | 0,338789 | 0,03616 | 9,369161 | < 2e-16 |
| ***Campylobacter jejuni / Tetracycline*** | | | | | |
| Intercept | | -0,574843 | 0,020942 | -27,449761 | < 2e-16 |

| | | | | | |
|---|---|---|---|---|---|
| Country | UK | 0 | | | |
| | USA | 0,329553 | 0,028736 | 11,468216 | < 2e-16 |
| Source | Human | 0 | | | |
| | Poultry | 0,00818 | 0,030118 | 0,271585 | 0.786 |
| | Farm animal | 0,179408 | 0,037583 | 4,773692 | 1.81e-06 |
| Year | Pre-2015 | 0 | | | |
| | 2015 | 0,183355 | 0,043425 | 4,222375 | 2.42e-05 |
| | 2016 | 0,450508 | 0,032462 | 13878 | < 2e-16 |
| | 2017 | 0,246668 | 0,036339 | 6,788023 | 1.14e-11 |
| | 2018 | 0,220927 | 0,033466 | 6,601617 | 4.07e-11 |

**Supplementary Figure S1.** Comparison of the proportion of UK and US *C. coli* samples resistant to aminoglycosides, macrolides, quinolones and tetracycline for individual years from 2001-2018.

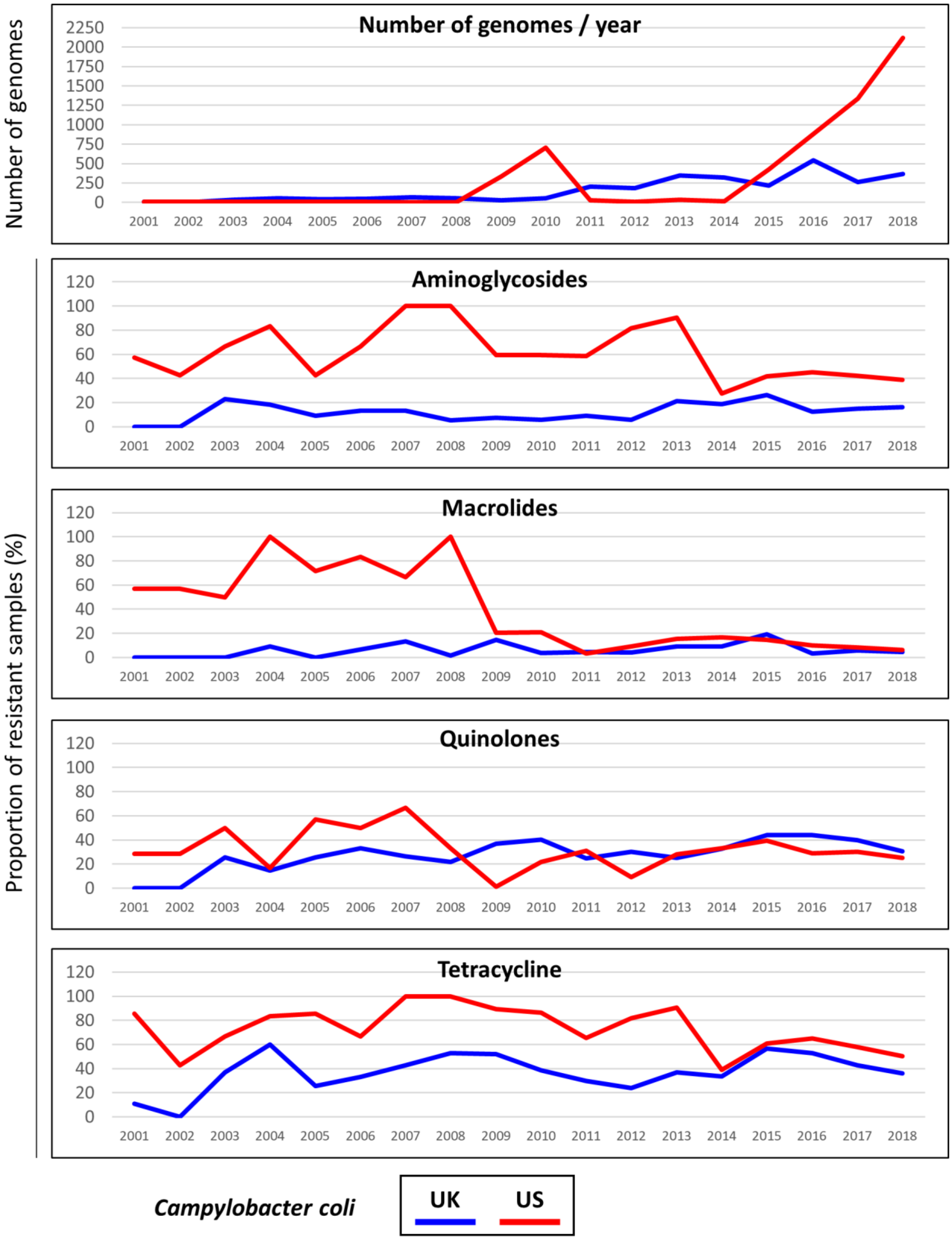

**Supplementary Figure S2.** Comparison of the proportion of UK and US *C. jejuni* samples resistant to aminoglycosides, macrolides, quinolones and tetracycline for individual years from 2001-2018.

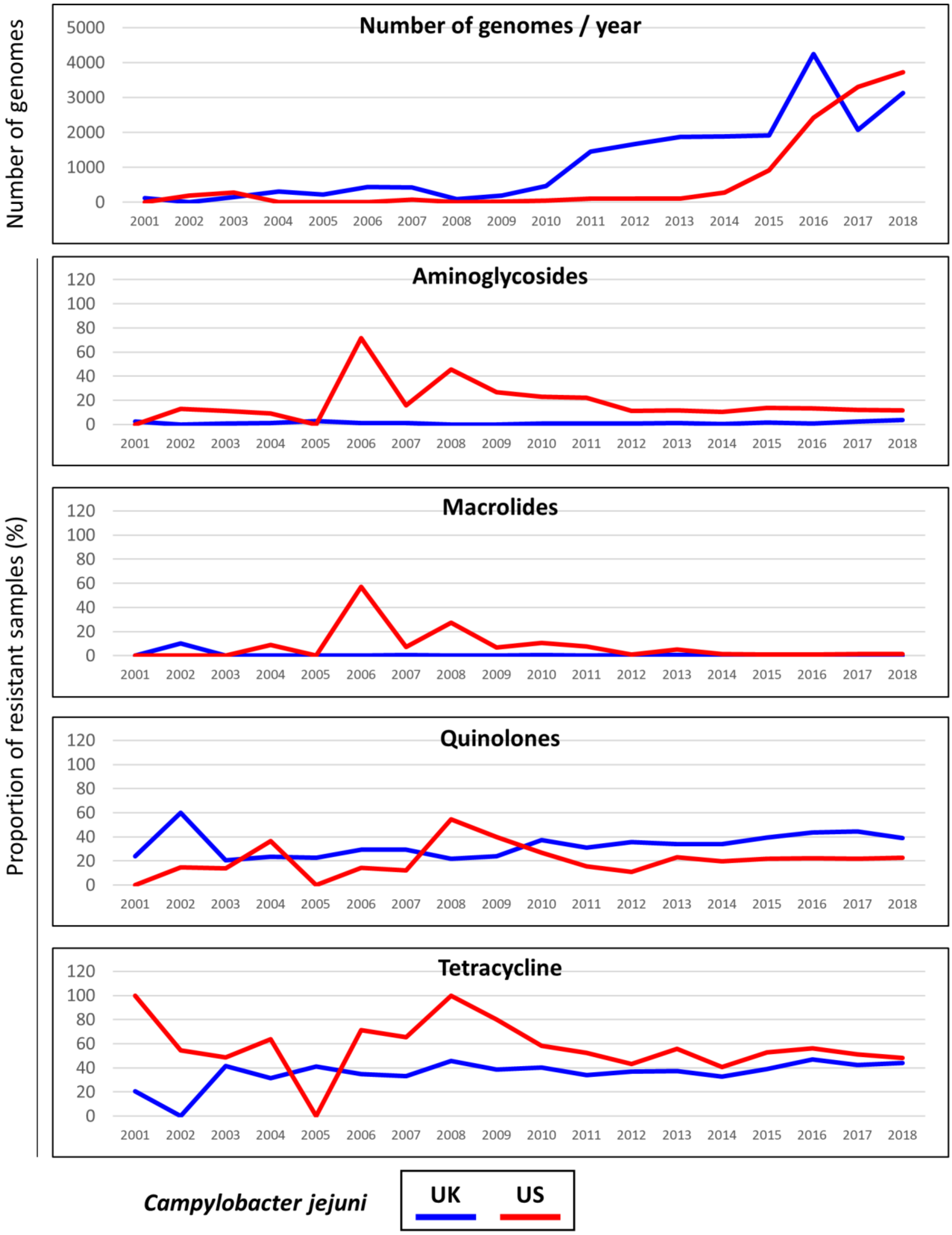
